# Metronomic doses and drug schematic combination response tested within microfluidic models for the treatment of breast cancer cells (JIMT-1)

**DOI:** 10.1101/2020.07.16.188748

**Authors:** Gustavo Rosero, Gisela Pattarone, Ana Peñaherera, Julia Felicitas Pilz, Joschka Bödecker, B Lerner, Roland Mertelsmann, M.S Perez, Marie Follo

## Abstract

Low-dose metronomic (LDM) chemotherapy is an alternative to conventional chemotherapy and is the most common use of low-dose levels of traditional chemotherapeutics in patients. The selection of patients, drug dosages and dosing intervals in LDM is empirical. In this study we systematically examined the schedule-dependent interaction of drugs on a breast cancer cell line (BCC) cultured in Lab on a Chip (LOC) microdevices. The LDM studies were combined with cell staining in order to better characterize different cell stages and modes of cell death, including caspase-dependent apoptosis, caspase-independent cell death and autophagy-dependent cell death. Microscope images were examined using the Fiji plugin Trainable Weka Segmentation to analyze cell area in 7500 images showing different types of cell death modes. Paclitaxel combined with LDM chemotherapy demonstrated a reduction in the area covered by live cells. In contrast, there was an induction of high levels of cell death due to caspase-dependent apoptosis. Furthermore, the microdevice used in this study is also an attractive alternative for staining cells in order to characterize and study BCC growth and development in situ.

## 1. Introduction

Discovering ways to fight or block malignant growth has been a driving force in cancer biology research over the past four decades^1^. Human breast cancer (BC) is the most common malignancy in women^2^. This type of cancer is comprised of phenotypically diverse populations of breast cancer cells (BCC)^3^. Paclitaxel has been demonstrated to have initial activity in BC^4^, and when combined with doxorubicin is the most active agent known to work against BC when using different schedules^5^. Low-dose metronomic (LDM) chemotherapy is a novel use of chemotherapy, and is defined as using conventional low doses with no prolonged drug-free periods^6^. Indeed, inter-tumor heterogeneity has been observed in breast carcinomas from various individuals^7^. In addition, tumor heterogenicity can influence biological behavior and therapeutic response^8^. Therefore, anticancer medications and tumor heterogeniciy can incite different modes of cell death, including caspase-dependent apoptosis, caspase-independent cell death, reproductive cell death (cell senescence), or cell death due to autophagy^9^. These non-standard pathways show the unexpected intricacy of molecular crosstalk within the cell death pathways, reflected in a diversity of phenotypes which have not yet been characterized in microfluidic models^10^. Microfluidics allow miniaturization of basic conventional biological or chemical laboratory processes. Lab-on-a-chip technology (LOC) has been well-accepted by biological and medical research communities as a promising tool for engineering the microenvironment at the cellular level^11^. Cell culture in microfluidic systems is not only able to provide suitable cell culture environments, it also permits furnishing cells with new media containing oxygen, carbon dioxide and supplements, while expelling metabolic products at a controlled flow rate ^1,12^. The unique characteristics of microfluidics in cell culture, such as fine geometric patterns, laminar liquid flow, and mixing purely based on diffusion, provides a diverse range of fluidic control strategies. A number of methods have recently been developed which enable modulation of the extracellular environment via dynamic control of the concentration of signal molecules and permit analyzes at the single cell level^(13,14,15,16,17,18,19)^. The characteristics of LOC devices make them especially suitable for cancer cell research.

Tumor cells showed increased viability, invasion and drug responses when they were seeded in LOC culture systems^20^. Conventional in vitro 2D assays have been widely used to evaluate the role of chemoattractants on cancer cell migration^21,22^. *Wang et al*. utilized a 2D platform simplifying the tumor microenvironment, not limited to cells growing in monolayers. They developed a microfluidic platform in order to stain cells inside of LOC devices^23,24^. In fact, staining cells inside microfluidic cultivation devices is a powerful tool to detect intracellular changes for live cell imaging. It allows the determination of changes in cell states such as distinguishing live or dead cells, lysed or senescent cells, and other dynamic changes in cell state^25,26,27,28^. For that reason, being able to develop new approaches for cell analyzes will allow this heterogeneity to be evaluated. Considering the benefits of LOC combined with the advantages of being able to use low numbers of cells and reagents, the present study used LOC microdevices for the study and characterization of biological behavior of breast cancer cells (JIMT-1) using different stainings to evaluate cell viability (LC), caspase-dependent programmed apoptosis(AP), autophagy (AG), and cell death (PI), inside of microchamber devices before and after the introduction of two different types of drug treatments applied *in vitro* with breast cancer cells.

## 2. Materials and Methods

### 2.1 Cell Culture

JIMT-1 cells ATCC 589 (DSMZ) were cultured in complete DMEM medium (Gibco), supplemented with fetal calf serum heat-inactivated (FBS) 10% (w/v) (Gibco), L-glutamine 2mmol·L^−1^ (Gibco), penicillin 100 units·mL^−1^, streptomycin 100 μg·mL (Gibco) at 37 °C in an incubator with 5% CO_2_. For microscopy assays, 20,000 cells per well were cultured in μ-Slide 8 well chambered polymer bottom coverslips (cat. no. 808126, Ibidi GmbH).

### 2.2 Cell death modes and characterization

Live/Dead Cell Imaging Kit (Sigma), Autophagy Cell Imaging Kit (CYTO-ID), Caspase-3 and −7 Cell Imaging Kit (Invitrogen), and Propidium iodide **(PI)** (Sigma-Aldrich) were used to identify the different cell death modes. For the Live/Dead **(LC)** assay, the viable cells are stained green, while the dead cells are stained red^29^. Autophagous cells in the autophagy (Figure 1) assay **(AG)** are stained green, and negative and positive controls were performed as recommended by the manufacturer’s instructions (Enzo ENZ-51031-K200)^30^. In the caspase-3 and −7 assay **(AP)**, the apoptotic cells are stained green^31^. In the **PI** assay, dead cells are stained red. The cells were loaded into the microfluidic devices, as described previously, and PI labeling, **LC, AG, AP**, were evaluated on the first, third, fourth and fifth days after treatment. We used phosphate buffered saline (PBS, HyClone) to wash the culture chambers in the models for 3 min. Cells were then incubated with each Imaging Kit for 30 min at 37 °C. Next, PBS was used to wash out the reagent for 5 min. Before examining the culture chambers under a fluorescent microscope, the cell culture chips were filled with fresh DMEM media. Finally, cell viability was measured by assessing the percentage of fluorescent cells in the cell cultures.

**Figure 1.-.**
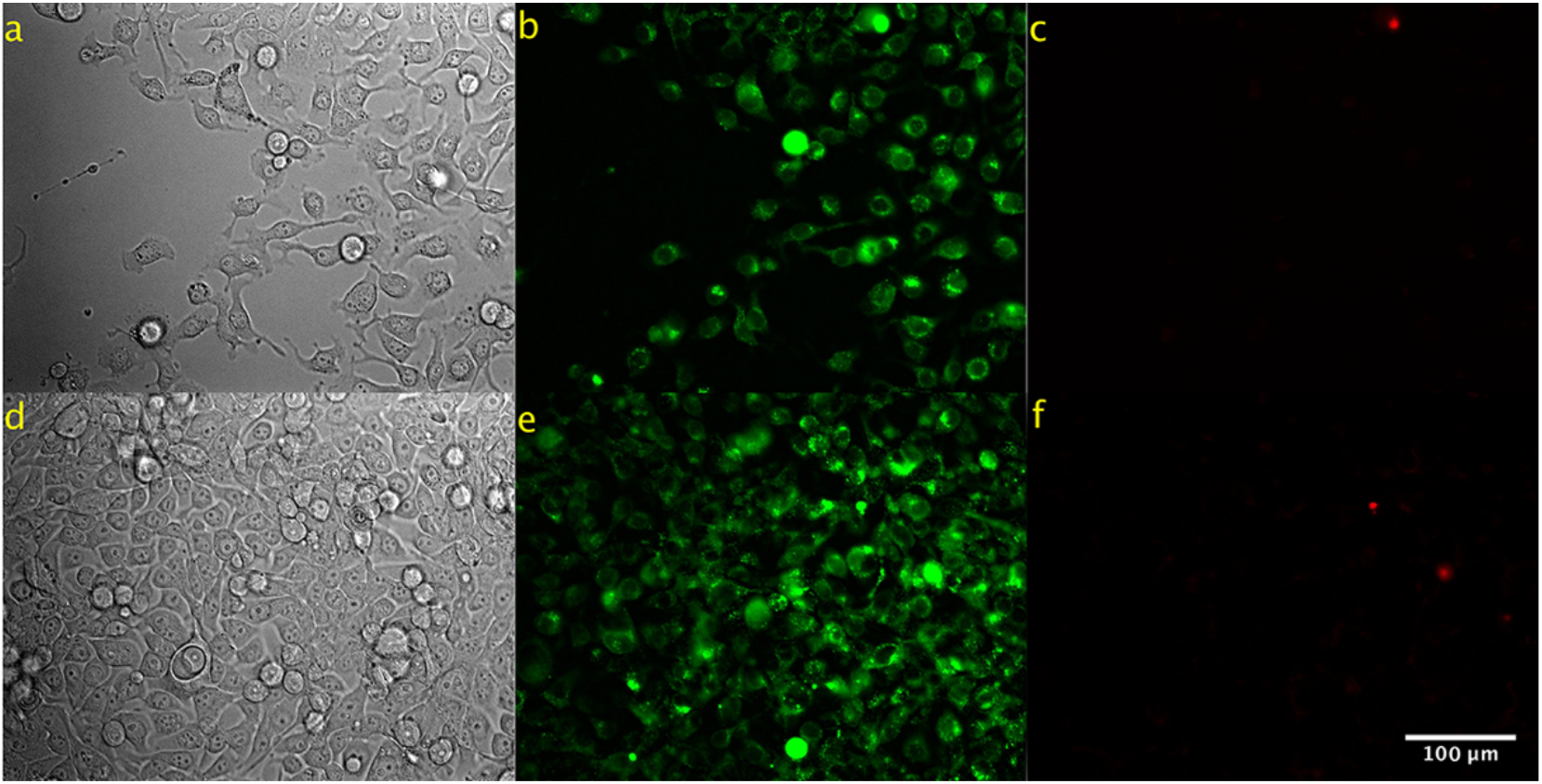
Representative microscopy images of autophagy and PI staining in breast cancer cells after three days of growing in two different wells in the microdevices. (a,d) Brightfield images of the culture. (b,e) Fluorescence images of cells after treatment with autophagy kit which preferentially labels autophagous vacuoles, shown here in green (c,f) Fluorescence images after treatment the cells with PI. Dead cells are shown in red.

### 2.3 Combined treatment of cells with Paclitaxel and doxorubicin and examination of drug effects

Paclitaxel and doxorubicin were added to the cells to test their effect, both alone and in combination (Sigma Aldrich)^34^. First, JIMT-1 cells were plated on the chambered coverslips and were grown for two days. After 24 h we replaced the medium with fresh culture medium mixed with doxorubicin at a concentration of 0.01 μM. After four hours in doxorubicin paclitaxel at a final concentration of 0.001 μM was added to the medium. The cells were cultured for a further 24 hours Live cell imaging and biological characterization of the cells were carried out using the different stainings as described above. Characterization and cell death mode analyzes were performed in order to evaluate the treatment efficiency. The first stage was after four hours of doxorubicin exposure and the second stage after doxorubicin and paclitaxel.

### 2.4 Metronomic Paclitaxel treatment and evaluation of drug effects

For the metronomic Paclitaxel tests, the effects of paclitaxel given at different time points were studied^35^. First, JIMT-1cells were plated onto the chambered slips as described above, after 24 hours the medium was replaced with fresh culture medium with paclitaxel at a concentration of 0.001 μM. The medium was then replaced with fresh culture medium containing paclitaxel at a concentration of 0.001 μM every 24 hours for the following five days. Biological characterizations using the different stainings were performed as described above.

### 2.5 Image processing and quantification

Brightfield and fluorescent images were obtained using an automated fluorescence microscope (Olympus ScanR High Content Screening Station) using the 20x UPLSAPO NA 0.75 objective. Cell images were classified according to cell viability (LC), programmed apoptosis (AP), autophagy (AG), or cell death (PI). In total, 7500 images were analyzed using the Fiji program (v.2.0.0) with the Trainable Weka Segmentation plugin (v.3.2.33): The Weka plugin within ImageJ is an open-source platform for biological-image analyzes that combines a collection of machine learning algorithms with a set of selected image features to produce pixel-based segmentations^36,37^. All of the experiments were performed in triplicate and the data are presented as the mean ± standard deviation (SD). A one-way analyzes of variance (ANOVA) and Student’s t-test were used for comparisons of each group. P-values less than 0.05 were considered statistically significant and are indicated with asterisks.

## 3. Results and Discussion

### 3.1 Tracking the growth of breast cancer cells within the cell culture chambers

Evaluation of the different schematics and the efficacy of therapeutic drugs is important. Rare publications have reported the effects of drugs in the characterization of breast cancer cells inside of microfluidic models. Chambered coverslips were used to track and evaluate the development of breast cancer cells over a period of five days. To better understand the development and characterization of cells within the slide chambers, the percentages of live, dead, autophagous, and apoptotic cells were analyzed over a period of five days. The results of growing cells under these different conditions can be observed in Figure 2.

**Figure 2.**
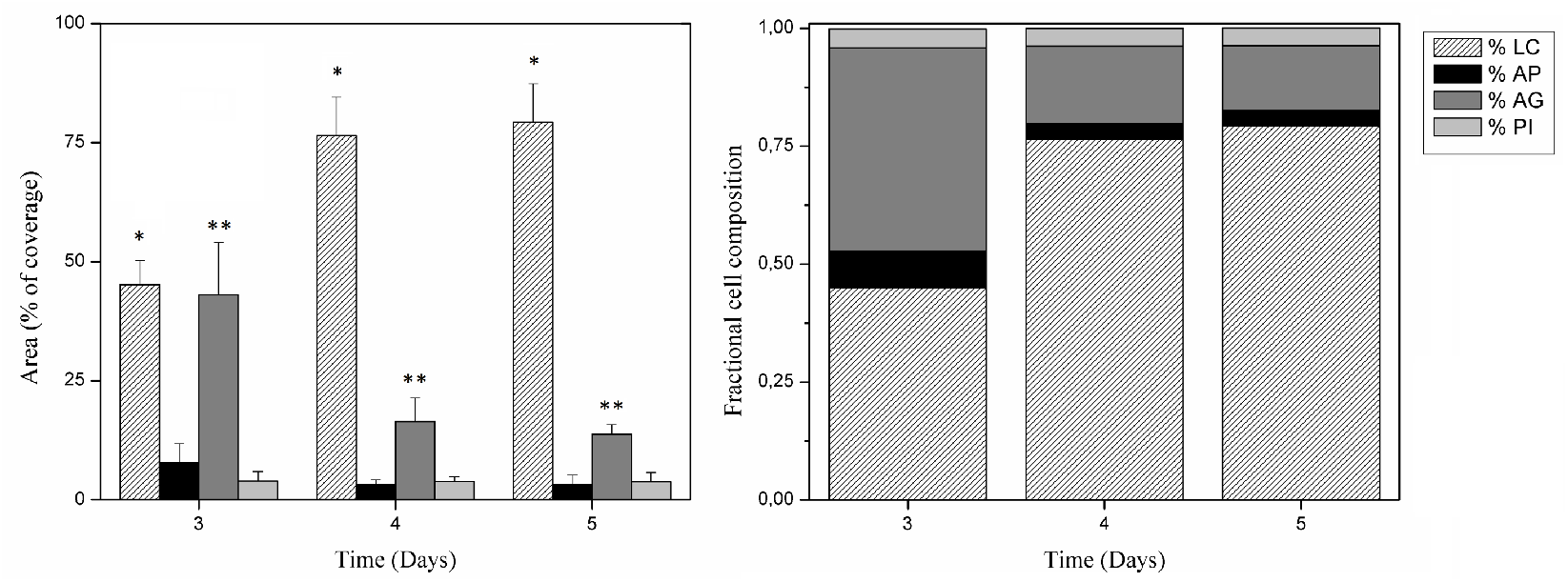
Cell distribution after characterization with stainings for 5 days. Right, stacked bar graph illustrating cell fraction composition per day over the five day period. Left, the surface area covered by live cells (**LC**) increased while the levels of the autophagy process decreased (**AG**) significantly. (*) and (**) indicate significant difference between days among the subgroups (p < 0.05).

Significantly differences in the percentages of **LC** and **AG** were observed over the five day time period. The percentage of live cells increased over the five day period (Figure 2 left). In contrast, cells in autophagy were at a high percentage of total cell area on the third day but had decreased by the fifth day. Notably, there is growth of the cells over the five day period, detectable when using the marker for live cells. The chambered coverslips were able to provide homogeneous conditions for keeping the cells viable in culture over this time period^38^.

### 3.2 Effect of Paclitaxel and doxorubicin drug schematics and the evaluation of LC, AG, PI and AP

There has been considerable interest in recent years in synergistic chemotherapeutic agents for the treatment of breast cancer^39,40^. Rare publications report the effect of drugs on the characterization of breast cancer cells within microfluidic model systems. On the one hand, previous clinical trials have shown benefits in combining DOX and PTX while treating breast cancer patients. However, side effects such as neutropenia and congestive heart failure have limited the dose administered to patients^41,42,43^. To analyze the effect of drugs on cells using the chambered coverslips, it is appropriate to keep the cell culture growing under homogeneous conditions^44^. First the effect of DOX was studied and the analyzes is described in Figure 3. Samples were incubated for four hours with DOX at a concentration of 0.01 μM. The level of apoptosis in the cells was estimated by quantifying the stained area relative to the entire cell population using the **AP** assay described above. Significant differences in apoptotic activity with respect to the control cells were found after four hours of exposure to DOX. The effect of DOX in the first four hours reduced the percentage in the area of living cells and produced an increase of apoptotic cells.

**Figure 3.**
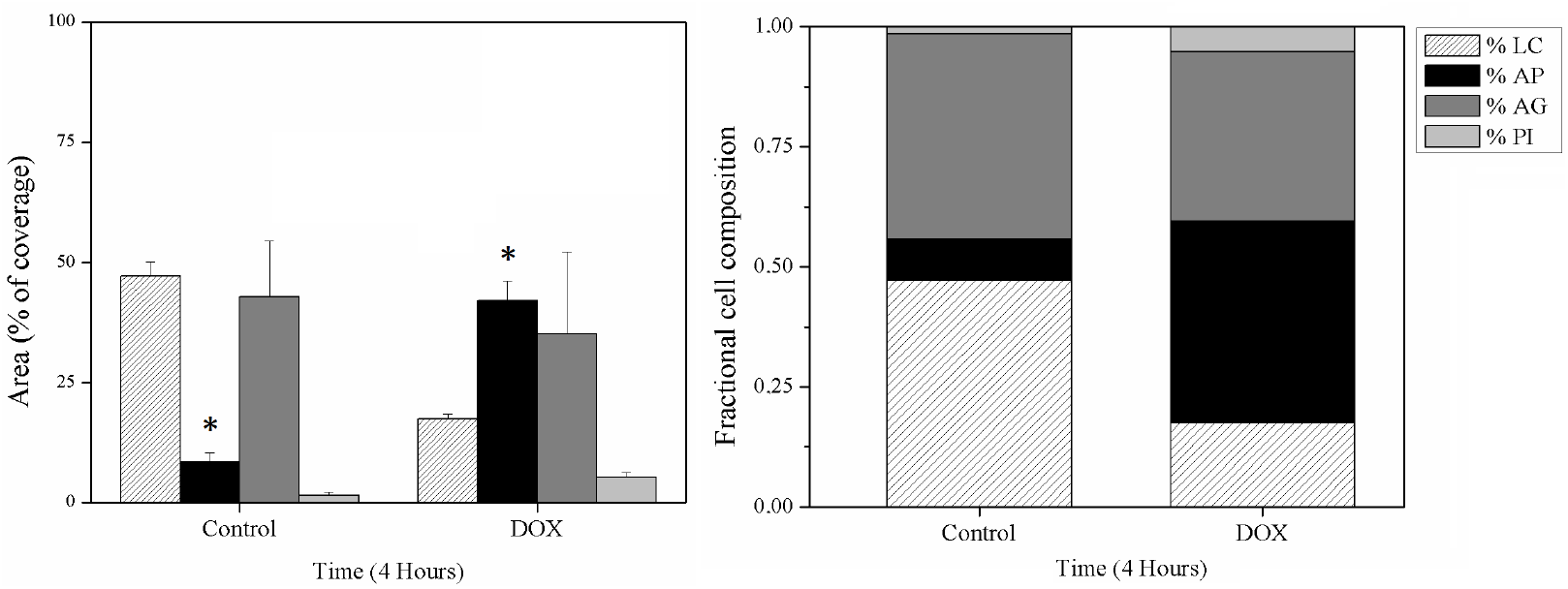
Doxorubicin effect on cells after four hours. Left, the apoptosis process (**AP**) increased significantly with respect to the control. The coverage area by Live cells (**LC**) decreased not significantly in comparison to the control. Right stacked bar graph shows the value of the results in cell fraction composition. (*) indicates significant difference with days among subgroups (p < 0.05).

The results obtained from the image analyzes can be seen in Figure 4. No significant differences were seen in LC, AP and AG, however the area stained for **PI** showed a slight but significant increase compared to the controls

**Figure 4.**
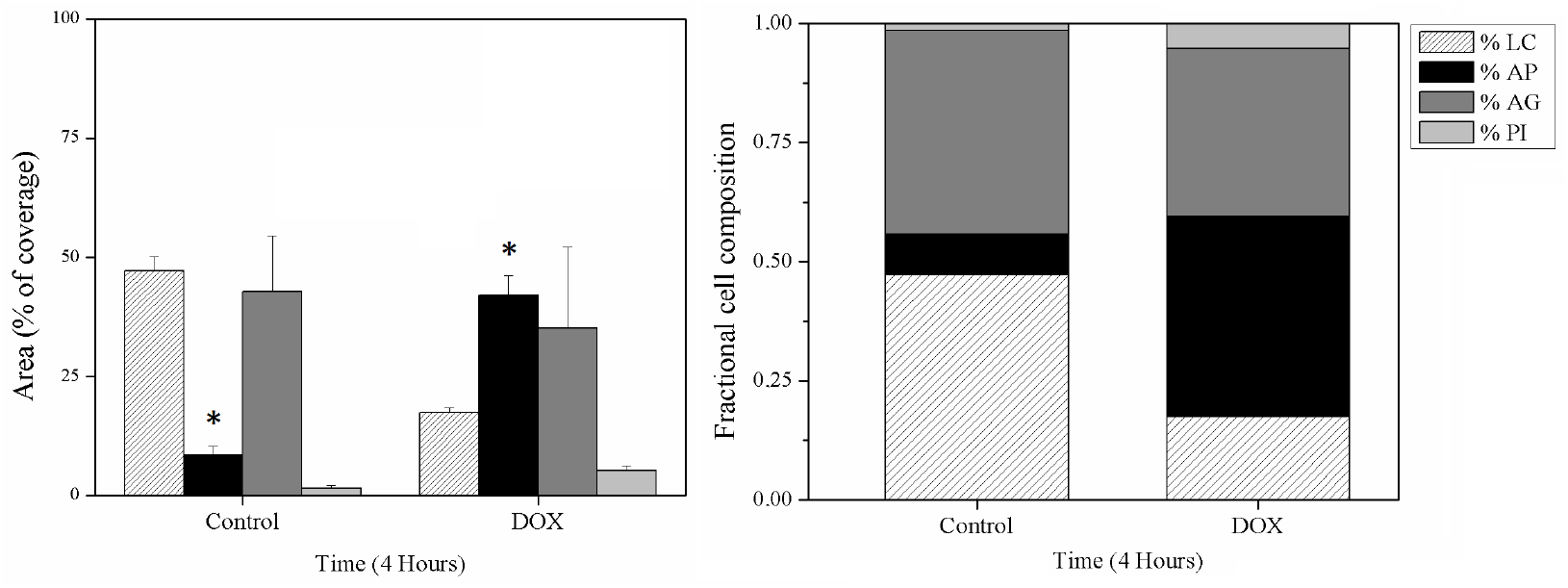
Doxorubicin combined with Paclitaxel. Bar graph denotes the effect of PTX after 24 hours on cells death modes. Propidium staining (**PI**) increased significantly with respect to the control. the area covered by live cells (**LC**) decreased not significantly in comparison to the control. Right stacked bar graph shows the value of the results in cell fraction composition per day along of four hours. (*) indicates significant difference with days among subgroups (p < 0.05).

Next, the effect of PTX combined with DOX is widely reported to have a deeply synergistic inhibitory effect on the growth of several breast cancer cell lines^46,47^. Our results (Figure 4) did not show further increases in the percentages of **AP** or of growth inhibition. This may be due to the fact that higher concentrations of PTX can actually have the opposite effect on cell growth, as has been reported earlier^48^. Numerous factors contribute to paclitaxel resistance within a given cell population, and these factors were found to be highly variable. Therefore, the exact mechanism its action remains unclear^48,49^.

### 3.3 Effect of metronomic treatment with Paclitaxel

The efficacy of paclitaxel was tested at very low concentrations over a period of five days. We observed that this drug had a great effect on LC (Figure 5), causing both a decrease in the area of **LC** and an increase in the cells undergoing **AP** with respect to the controls (Figure 5). The area of cells with **AG** and **PI** remained almost constant over the same time periods.

**Figure 5.**
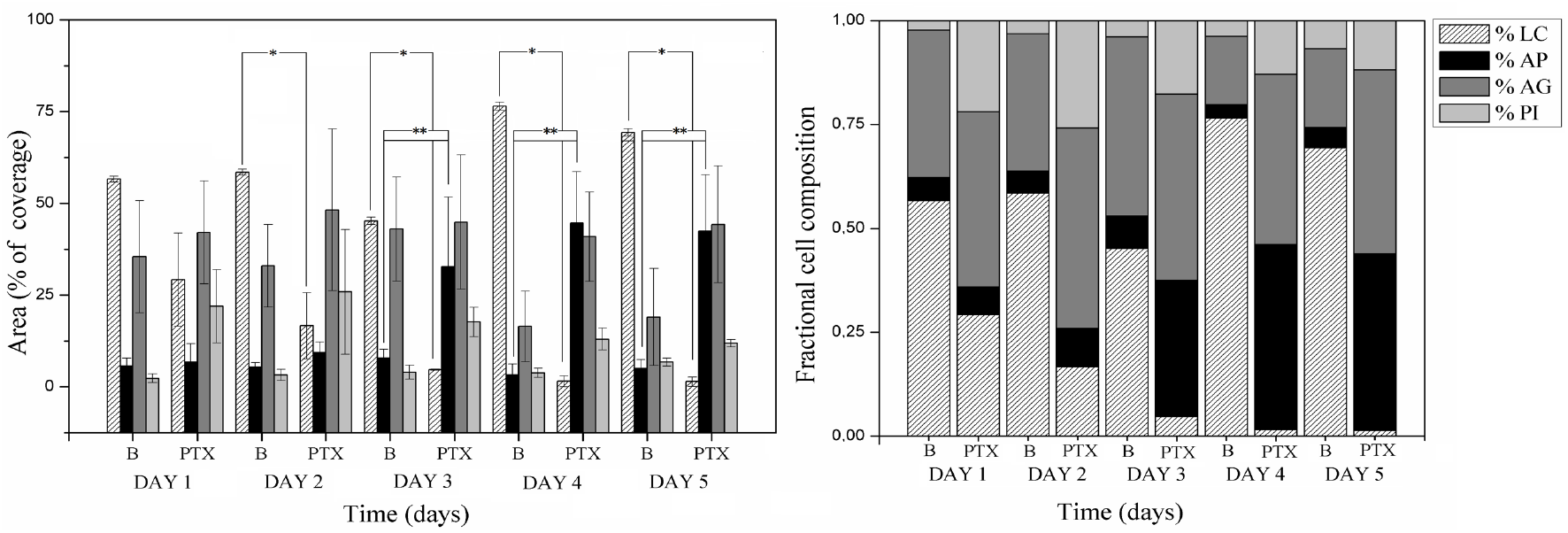
Paclitaxel (PTX) effect compared with control (B) over 5 days at low PTX concentration. Bar graph denotes the value of percentage of the area covered by cells for each of the death modes studied. Left, the apoptosis process (**AP**) increased significantly in comparison to the control, and the area coved by live cells (**LC**) decreased significantly with respect to the control. Right, stacked bar graph shows the value of the results in cell fraction composition per day a period of five days. (**) and (*) indicates significant difference between days among subgroups (p < 0.05).

Metronomic treatment (LDM) with PTX was found to lead to a higher percentage of AP cells and a considerable decrease in the overall percentage of live cells. LDM therapy was widely reported in recent years in clinical trials and in *in vitro* studies using PTX at very low concentrations^50,51,52^. PTX in LDM (figure 7B) was found to induce **AP** and inhibit cell growth, suggesting that it induces mitotic cell death by disrupting microtubule dynamics and activating the spindle-assembly checkpoint (SAC)^53^.

In addition, one of the proapoptotic pathways evaluated was **AG**. Without drugs we observed that the percentage of autophagous cells did not decrease over the period of several days studied but rather remained constant. Apparently, this mechanism can provide extra energy and resources to the cancer cells, allowing their survival in microfluidic devices^54^. The percentage of AG cells remained constant under drug treatment in our experiments. These results are in agreement with the fact that cancer cells exposed to chemotherapy can significantly impact the induction of proapoptotic pathways^45^.

In summary, it was possible to characterize the growth of breast cancer cells within cell chambers and characterize them following different treatments. PTX with LDM was found to be the best treatment to induce apoptosis in these cells in LOC. Furthermore, the **AG** pathway showed an unexpected mechanism for these cells to be able to adapt to growth in the LOC as well as to the drug treatment.

### Conclusions

These results confirm the usefulness of LOC as an effective method for drug dosage quantification of live, dead, autophagous, and apoptotic cells. LOC allows comparing two different treatments for breast cancer cells. The metronomic treatment (LDM) reported a considerable decrease in the overall percentage of live cells and increase in the cells undergoing **AP** with respect to the controls. In conclusion, LOC devices have not only the already known advantages, they are also a powerful tool since they require only slight amounts of reagents and can be used under specific culture parameters. LOC devices could be considered as a novel technology to be used as a complement or replacement of traditional studies on metronomic doses of drugs with different treatments and schematics.

## Acknowledgements

We would like to thank Juan Pablo Yánez for his kindness and for helping the authors during their stay in Freiburg. Also, to Hans Hoch and his family for their support during the research. Furthermore, the authors feel thankful to the Biothera Foundation for giving the authors the opportunity to visit the Lighthouse Core Facility lab and to the members of the lab for helping us every time, especially to our friend Klaus Geiger who will always live in our memories.

## References

1. Hanahan, D. & Weinberg, R. A. Hallmarks of Cancer: The Next Generation. Cell 144, 646–674 (2011).

2. Arabsalmani, M. et al. Incidence and mortality of kidney cancers, and human development index in Asia; a matter of concern. J. Nephropathol. 6, 30–42 (2017).

3. Al-Hajj, M., Wicha, M. S., Benito-Hernandez, A., Morrison, S. J. & Clarke, M. F. Prospective identification of tumorigenic breast cancer cells. Proc. Natl. Acad. Sci. 100, 3983 (2003).

4. Reichman, B. S. et al. Paclitaxel and recombinant human granulocyte colonystimulating factor as initial chemotherapy for metastatic breast cancer. J. Clin. Oncol. 11, 1943–1951 (1993).

5. Pazos, C. et al. Phase II of doxorubicin/taxol in metastatic breast cancer. Breast Cancer Res. Treat. 55, 91–96 (1999).

6. Lien, K., Georgsdottir, S., Sivanathan, L., Chan, K. & Emmenegger, U. Low-dose metronomic chemotherapy: A systematic literature analysis. Eur. J. Cancer 49, 3387–3395 (2013).

7. Turashvili, G. & Brogi, E. Tumor Heterogeneity in Breast Cancer. Front. Med. 4, 227–227 (2017).

8. Belgorosky, D. et al. Analysis of tumoral spheres growing in a multichamber microfluidic device. J. Cell. Physiol. 233, 6327–6336 (2018).

9. Leist, M. & Jäättelä, M. Four deaths and a funeral: from caspases to alternative mechanisms. Nat. Rev. Mol. Cell Biol. 2, 589 (2001).

10. Wlodkowic, D., Skommer, J. & Darzynkiewicz, Z. Cytometry in cell necrobiology revisited. Recent advances and new vistas. Cytometry A 77A, 591–606 (2010).

11. Xu, Z. et al. Application of a microfluidic chip-based 3D co-culture to test drug sensitivity for individualized treatment of lung cancer. Biomaterials 34, 4109–4117 (2013).

12. Leclerc, E., Sakai, Y. & Fujii, T. Cell Culture in 3-Dimensional Microfluidic Structure of PDMS (polydimethylsiloxane). Biomed. Microdevices 5, 109–114 (2003).

13. Zare, R. N. & Kim, S. Microfluidic Platforms for Single-Cell Analysis. Annu. Rev. Biomed. Eng. 12, 187–201 (2010).

14. Cheong, R., Wang, C. J. & Levchenko, A. High Content Cell Screening in a Microfluidic Device. Mol. Amp Cell. Proteomics 8, 433 (2009).

15. King, K. R., Wang, S., Jayaraman, A., Yarmush, M. L. & Toner, M. Microfluidic flow-encoded switching for parallel control of dynamic cellular microenvironments. Lab. Chip 8, 107–116 (2008).

16. Cheong, R., Wang, C. J. & Levchenko, A. Using a microfluidic device for high-content analysis of cell signaling. Sci. Signal. 2, pl2–pl2 (2009).

17. Lee, P. J., Gaige, T. A. & Hung, P. J. Dynamic cell culture: a microfluidic function generator for live cell microscopy. Lab. Chip 9, 164–166 (2009).

18. Eriksson, E. et al. A microfluidic device for reversible environmental changes around single cells using optical tweezers for cell selection and positioning. Lab. Chip 10, 617–625 (2010).

19. Kim, S.-J., Yokokawa, R., Lesher-Perez, S. C. & Takayama, S. Constant flow-driven microfluidic oscillator for different duty cycles. Anal. Chem. 84, 1152–1156 (2012).

20. Liu, T. et al. A microfluidic device for characterizing the invasion of cancer cells in 3-D matrix. ELECTROPHORESIS 30, 4285–4291 (2009).

21. Wang, S.-J., Saadi, W., Lin, F., Minh-Canh Nguyen, C. & Li Jeon, N. Differential effects of EGF gradient profiles on MDA-MB-231 breast cancer cell chemotaxis. Exp. Cell Res. 300, 180–189 (2004).

22. Saadi, W., Wang, S.-J., Lin, F. & Jeon, N. L. A parallel-gradient microfluidic chamber for quantitative analysis of breast cancer cell chemotaxis. Biomed. Microdevices 8, 109–118 (2006).

23. Wang, C., Lu, H. & Alexander Schwartz, M. A novel in vitro flow system for changing flow direction on endothelial cells. J. Biomech. 45, 1212–1218 (2012).

24. Toh, Y.-C., Raja, A., Yu, H. & van Noort, D. A 3D Microfluidic Model to Recapitulate Cancer Cell Migration and Invasion. Bioeng. Basel Switz. 5, 29 (2018).

25. Krämer, C. E. M. et al. Non-Invasive Microbial Metabolic Activity Sensing at Single Cell Level by Perfusion of Calcein Acetoxymethyl Ester. PLOS ONE 10, e0141768 (2015).

26. Hoshi, H., O’Brien, J. & Mills, S. L. A novel fluorescent tracer for visualizing coupled cells in neural circuits of living tissue. J. Histochem. Cytochem. Off. J. Histochem. Soc. 54, 1169–1176 (2006).

27. Byrd, T. F., 4th et al. The microfluidic multitrap nanophysiometer for hematologic cancer cell characterization reveals temporal sensitivity of the calcein-AM efflux assay. Sci. Rep. 4, 5117–5117 (2014).

28. Krämer, C. E. M. et al. Non-Invasive Microbial Metabolic Activity Sensing at Single Cell Level by Perfusion of Calcein Acetoxymethyl Ester. PLOS ONE 10, e0141768 (2015).

29. Zhang, J., Wu, J., Li, H., Chen, Q. & Lin, J.-M. An in vitro liver model on microfluidic device for analysis of capecitabine metabolite using mass spectrometer as detector. Biosens. Bioelectron. 68, 322–328 (2015).

30. Warenius, H. M. et al. Selective anticancer activity of a hexapeptide with sequence homology to a non-kinase domain of Cyclin Dependent Kinase 4. Mol. Cancer 10, 72 (2011).

31. Utsumi, F. et al. Effect of Indirect Nonequilibrium Atmospheric Pressure Plasma on Anti-Proliferative Activity against Chronic Chemo-Resistant Ovarian Cancer Cells In Vitro and In Vivo. PLOS ONE 8, e81576 (2013).

32. Utsumi, F. et al. Effect of Indirect Nonequilibrium Atmospheric Pressure Plasma on Anti-Proliferative Activity against Chronic Chemo-Resistant Ovarian Cancer Cells In Vitro and In Vivo. PLOS ONE 8, e81576 (2013).

33. Utsumi, F. et al. Effect of Indirect Nonequilibrium Atmospheric Pressure Plasma on Anti-Proliferative Activity against Chronic Chemo-Resistant Ovarian Cancer Cells In Vitro and In Vivo. PLOS ONE 8, e81576 (2013).

34. Holmes, F. A. et al. Sequence-dependent alteration of doxorubicin pharmacokinetics by paclitaxel in a phase I study of paclitaxel and doxorubicin in patients with metastatic breast cancer. J. Clin. Oncol. 14, 2713–2721 (1996).

35. Lien, K., Georgsdottir, S., Sivanathan, L., Chan, K. & Emmenegger, U. Low-dose metronomic chemotherapy: A systematic literature analysis. Eur. J. Cancer 49, 3387–3395 (2013).

36. Schindelin, J. et al. Fiji: an open-source platform for biological-image analysis. Nat. Methods 9, 676 (2012).

37. Arganda-Carreras, I. et al. Trainable Weka Segmentation: a machine learning tool for microscopy pixel classification. Bioinformatics 33, 2424–2426 (2017).

38. Song, J. W. et al. Microfluidic Endothelium for Studying the Intravascular Adhesion of Metastatic Breast Cancer Cells. PLOS ONE 4, e5756 (2009).

39. Turashvili, G. & Brogi, E. Tumor Heterogeneity in Breast Cancer. Front. Med. 4, 227–227 (2017).

40. Leist, M. & Jäättelä, M. Four deaths and a funeral: from caspases to alternative mechanisms. Nat. Rev. Mol. Cell Biol. 2, 589 (2001).

41. Holmes, F. A. et al. Paclitaxel by 24-hour infusion with doxorubicin by 48-hour infusion as initial therapy for metastatic breast cancer: Phase I results*. Ann. Oncol. 10, 403–411 (1999).

42. Fisherman, J. S. et al. Phase I/II study of 72-hour infusional paclitaxel and doxorubicin with granulocyte colony-stimulating factor in patients with metastatic breast cancer. J. Clin. Oncol. 14, 774–782 (1996).

43. Perez, E. A. Doxorubicin and Paclitaxel in the Treatment of Advanced Breast Cancer: Efficacy and Cardiac Considerations. Cancer Invest. 19, 155–164 (2001).

44. Song, J. W. et al. Microfluidic endothelium for studying the intravascular adhesion of metastatic breast cancer cells. PloS One 4, e5756–e5756 (2009).

45. Lee, M. J. et al. Sequential Application of Anticancer Drugs Enhances Cell Death by Rewiring Apoptotic Signaling Networks. Cell 149, 780–794 (2012).

46. Wang, L. et al. De-repression of the p21 promoter in prostate cancer cells by an isothiocyanate via inhibition of HDACs and c-Myc. Int. J. Oncol. 33, 375–80 (2008).

47. Wang, L. G. et al. Dual action on promoter demethylation and chromatin by an isothiocyanate restored GSTP1 silenced in prostate cancer. Mol. Carcinog. 46, 24–31 (2007).

48. Liebmann, J. E. et al. Cytotoxic studies of paclitaxel (Taxol) in human tumour cell lines. Br. J. Cancer 68, 1104–1109 (1993).

49. Huang, C. et al. FoxM1 Induced Paclitaxel Resistance via Activation of FoxM1/PHB1/RAF-MEK-ERK Pathway and Enhancement of ABCA2 Transporter. Mol. Ther. - Oncolytics 14, (2019).

50. Lien, K., Georgsdottir, S., Sivanathan, L., Chan, K. & Emmenegger, U. Low-dose metronomic chemotherapy: A systematic literature analysis. Eur. J. Cancer 49, 3387–3395 (2013).

51. Souto, M., Shimada, A., Barbosa, C. C., Cruz Abrahao, M. & Katz, A. Low-dose metronomic chemotherapy in metastatic breast cancer: A retrospective analysis of 40 patients. J. Clin. Oncol. 33, e11567–e11567 (2015).

52. Jiang, H. et al. Low-Dose Metronomic Paclitaxel Chemotherapy Suppresses Breast Tumors and Metastases in Mice. Cancer Invest. 28, 74–84 (2010).

53. Musacchio, A. & Salmon, E. D. The spindle-assembly checkpoint in space and time. Nat. Rev. Mol. Cell Biol. 8, 379–393 (2007).

54. Huang, Z., Zhou, L., Chen, Z., Nice, E. & Huang, C. Stress Management by Autophagy: Implications for Chemoresistance. Int. J. Cancer J. Int. Cancer 139, (2016).

